# The three-dimensional anatomy and dorsoventral asymmetry of the mature *Marchantia polymorpha* meristem develops from a symmetrical gemma meristem

**DOI:** 10.1101/2024.08.23.609409

**Authors:** Victoria Spencer, Eva-Sophie Wallner, Katharina Jandrasits, Natalie Edelbacher, Magdalena Mosiolek, Liam Dolan

## Abstract

Meristems are three-dimensional generative structures that maintain a population of stem cells whilst producing new organs and tissues. Meristems develop in all land plants, however we know relatively little about the spatial and temporal regulation of meristem structure in lineages such as the bryophytes. Here we describe the three-dimensional anatomy of the meristem during the development of the liverwort, *Marchantia polymorpha.* Using optical reconstructions of the frontal, sagittal and transverse planes through the mature meristem, we show that the apical stem cell is sub-apical, ventral, and located in the outer cell layer. The anatomy of the mature meristem is therefore asymmetrical in the dorsoventral axis, which is reflected by the domain specific protein localisation of Marchantia Class III and Class IV Homeodomain-Leucine-Zippers (MpC3HDZ and MpC4HDZ) and promoter activity of Mp*YUCCA2*. The dorsoventral asymmetry that defines the mature meristem is absent in the juvenile meristems of the asexual propagules known as gemmae. We discovered that anatomical dorsoventral asymmetry of the meristem forms after two days of gemmaling growth and is accompanied by expression of the dorsal identity reporter, MpC3HDZ. We conclude that the gemma meristem is in a state of arrested development and undergoes anatomical rearrangement to develop the three-dimensional meristem structure of the mature plant.

## INTRODUCTION

The indeterminate growth of land plants is dependent on the maintenance of generative centres known as meristems. Meristems are three-dimensional structures that contain stem cells and their immediate daughter cells (collectively termed the stem cell niche), as well as an undifferentiated, proliferative zone that rapidly divides to produce the tissues of the plant body (Eshed Williams, 2021). The anatomy and molecular regulation of flowering plant meristems are well described. Both the shoot apical meristem and root apical meristem are dome-shaped and produce organs such as leaves and lateral roots behind the stem cell niche. Meristem anatomy is maintained in a steady state by the spatial restriction of gene expression to discrete domains (Kitagawa & Jackson, 2019; Yamoune et al., 2021). This steady state is established de novo during meristem formation in the embryo and is modified during developmental transitions such as reproduction (Barton, 2010; Bertran et al., 2024). Meristem architecture is therefore dynamically regulated through the plant lifecycle.

All land plants produce meristems, however our understanding of how meristem architecture is regulated in non-flowering plants such as bryophytes is lacking. (Arnoux-Courseaux & Coudert, 2024). Bryophytes have a haploid dominant phase (known as the gametophyte), and it is hypothesised that these meristems evolved independently to the meristems of the diploid dominant phase of flowering plants (the sporophyte) (Fouracre & Harrison, 2022). Flowering plants and gymnosperms typically have a multicellular stem cell niche, whilst gametophyte bryophyte meristems often contain one or two stem cells known as apical cells. The shape and number of cutting faces of the apical cell varies in different lineages (Shimamura, 2016). For example, the apical cell of simple thalloid liverworts is typically lenticular with two cutting faces, whilst the apical cell of complex thalloid liverworts such as the Marchantiales is typically wedge-shaped with four cutting faces (Douin, 1925; Leitgeb, 1881; Shimamura, 2016). It is unclear how the spatio-temporal regulation of meristem architecture in each land plant lineage compares.

Molecular regulators of haploid meristem maintenance have been identified in bryophytes such as the liverwort *Marchantia polymorpha*. For example, CLAVATA LRR-RLK receptors and their CLAVATA3/Endosperm surrounding region-related (CLE) peptide ligands are conserved in *M. polymorpha* (Hirakawa et al., 2020; Takahashi et al., 2021). However, MpCLV1-MpCLE2 signalling promotes stem cell proliferation in the Marchantia meristem, in contrast to CLAVATA1/2-CLV3 signalling in flowering plants which inhibits stem cell proliferation (Hirakawa, 2022; Hirakawa et al., 2020). Other identified regulators include Mp*JINGASA* (*MpJIN*) (Takahashi et al., 2023); Mp*PLETHORA* (*MpPLT*) (Fu et al., 2024); Mp*AINTEGUMENTA* (Mp*ANT*) (Liu et al., 2024); and Mp*TDIF RECEPTOR* (Mp*TDR*) (Hirakawa et al., 2019). These studies suggest that some of the same genes control meristem function in *M. polymorpha* and flowering plants.

Our understanding of the genetic regulation of the Marchantia meristem has progressed significantly in recent years. However, current expression data in *M. polymorpha* meristems lack cellular resolution in three-dimensions and focus primarily on the early development of young meristems in vegetative propagules known as gemmae. To understand the spatio-temporal regulation of the networks that control *M. polymorpha* meristem development, a 3D framework of meristem anatomy is needed. Here we provide a description of the cellular organisation of the *M. polymorpha* meristem and show that the anatomy and gene expression is distinct in each plane and is dorsoventrally polarised. Inactive gemmae lack dorsoventral asymmetry, which becomes progressively established during the first 3 days of gemmaling growth. We conclude that the mature gemma meristem is arrested and undergoes substantial anatomical development to produce the steady-state meristem of the mature plant.

## RESULTS

The *M. polymorpha* thallus is flat and bifurcating when grown in white light. Growth and morphogenesis occur at the tips of the thallus in notches between two lobes of differentiated tissues. In the notch, an apical cell acts as a self-renewing stem cell, which together with its immediate daughter cells constitute the stem cell niche. This stem cell niche produces undifferentiated proliferating cells that form the surface and internal tissues of the thallus. Collectively, the stem cell niche and the zone of undifferentiated, proliferating cells constitute the meristem (Arnoux-Courseaux & Coudert, 2024; Kohchi et al., 2021; Shimamura, 2016).

To define the three-dimensional cellular architecture of the mature meristem, wild-type plants were grown from gemmae for 4 weeks in white light before fixation and clearing. The meristem was imaged in each of the three anatomical planes. These include the frontal plane, which bisects the dorsal and ventral sections of the plant body; the sagittal plane, which bisects the left and right sections; and the transverse plane, which bisects the apical and basal sections of the plant (Fig. 1A). The predicted apical cell was identified based on its geometry and its position in the central axis of the notch.

**Figure 1:**
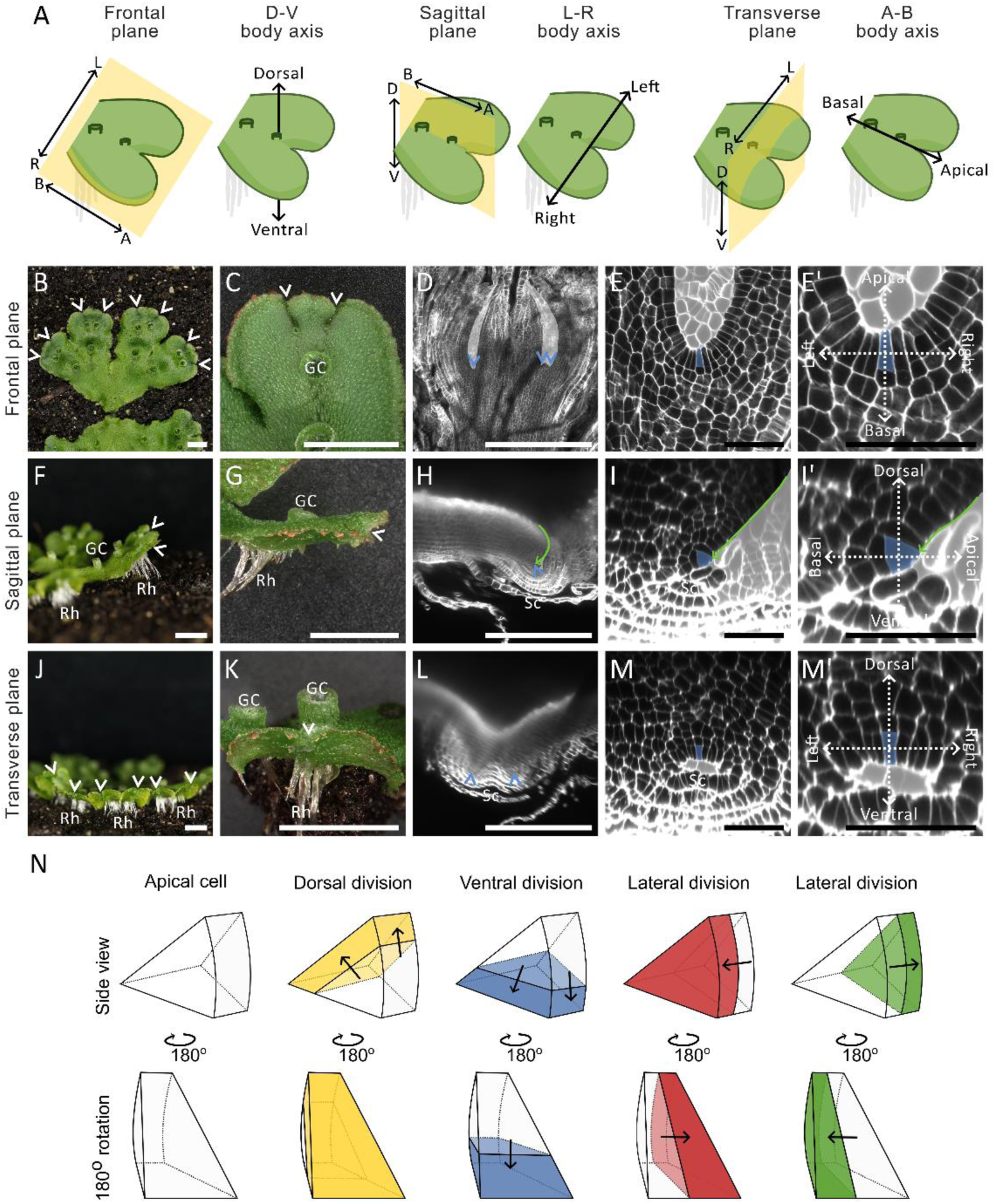
The anatomy of the mature *M. polymorpha* meristem is bilaterally symmetrical. **(A)** Schematic representation of the anatomical planes and body axes of the mature Tak-1 *Marchantia polymorpha* meristem. The frontal plane separates the dorsal (D) and ventral (V) sides, the sagittal plane separates the left (L) and right (R) sides, and the transverse plane separates the apical (A) and basal (B) sides of the plant body. The meristem is positioned where the three planes intersect. (**B**-**E**’) Frontal views of the *M. polymorpha* plant. (**B**) The bifurcating plant body grows horizontally along the substrate. (**C**) Meristems are located at the tip of the plant in notches (white arrowheads). Gemma cups (GC) are visible on the thallus surface. (**D**) Confocal microscopy image showing the stem cell niche (blue arrow heads) within the notch. (**E**-**E’**) Confocal microscopy image showing that the predicted apical cell (blue) is located at the base of the notch in the outer cell layer. E’ is a close up of E. The apical-basal and left-right body axes are marked. (**F**-**I**’) Sagittal views of the *M. polymorpha* plant. (**F**) Rhizoids (Rh) grow from the ventral surface and gemma cups (GC) protrude upwards from the dorsal surface of the thallus. (**G**) Meristems are located at the tip of the thallus, but the notch (white arrowhead) is not visible from this orientation. (**H**) Lightsheet microscopy image showing the stem cell niche (blue arrowhead) at the apex of the thallus (green arrow). The stem cell niche is surrounded by scales (Sc). (**I**-**I’**) Confocal microscopy image showing the apical cell (blue) is positioned sub-apically and on the ventral side of the thallus apex. I’ is a close up of I. The dorsoventral and apical-basal body axes are marked. (**J**-**M’**) Transverse views of the *M. polymorpha* plant. (**J**) Rhizoids (Rh) form from the ventral surface. The heart-shaped notch (white arrowhead) is evident due to the slight upwards bending of the thallus. (**K**) The meristem is in the base of the notch (white arrowhead) and is flanked by the thallus lobes. Gemma cups (GC) protrude from the dorsal thallus surface behind the notch. (**L**) Light sheet microscopy image showing the stem cell niche (blue arrowheads) on the ventral surface of the thallus, surrounded by scales. (**M**-**M’**) Confocal microscopy image showing the ventrally localised apical cell (blue). M’ is a close up of M. The dorsoventral and left-right body axes are marked. In all confocal images, the space surrounding the meristem is marked in pale grey for clarity. (**N**) The stem cells in the meristem are wedge-shaped in three dimensions. The stem cells divide in four directions to produce daughter cells. Together the stem cells and their immediate daughter cells form the stem cell niche. Scale bars: B-C, F-G, J-K= 5 mm; D, H, L= 500 µm; E-E’ I-I’, M-M’ = 50 µm.

In the frontal plane, the stem cell niche of the mature meristem was positioned at the base of the notch (Fig. 1B-D). The apical cell was within the outer cell layer and appeared trapezoidal in this orientation (Fig. 1E-E’). The tissue morphology was identical on the left and right of the stem cell niche. In the sagittal plane, the mature *M. polymorpha* thallus comprised many tissue layers. The outermost layers of the dorsal and ventral surfaces are morphologically distinct. The roof of air chambers, and gemma cups developed on the dorsal surface, while rhizoids and scales formed on the ventral surface (Fig. 1F-G). The apical cell was located sub-apically in the outer cell layer, behind the apex of the thallus (green arrow, Fig. 1H-I’). The outer cell layer on the apical side of the stem cell niche was curved and morphologically distinct from the parallel dorsal and ventral outer cell layers on the basal side of the stem cell niche. The apical cell was triangular in the sagittal plane and surrounded by tissue protrusions (scales) on the ventral surface of the meristem (Fig. 1I). The ventral side of the meristem was therefore morphologically distinct from the dorsal side of the meristem.

In the transverse plane, the stem cell niche of the mature meristem was located on the ventral side of the thallus, above the scales (Fig. 1J-L). The stem cell niche was flanked by the left and right thallus lobes and the apical cell was trapezoidal in this plane (Fig. 1M-M’). The tissue morphology was identical on the left and right of the meristem.

Imaging the meristem in three planes showed that the apical cell within the stem cell niche was trapezoid in the frontal and transverse planes, and triangular in the sagittal plane. Consistent with published results, the apical cell was therefore wedge-shaped and likely divided in four directions based on its geometry (Fig. 1N). Our analysis indicated that the cellular anatomy of the mature meristem was identical on the left and right, but different on the dorsal and ventral sides, and different on the apical and basal sides of the meristem. We conclude that the three-dimensional mature meristem has only one plane of symmetry-the sagittal plane- and is therefore bilaterally symmetrical.

The maintenance of meristem structure in flowering plants involves the spatial restriction of gene expression to distinct domains within the meristem. Our analysis of the mature meristem showed that the apical cell is located on the ventral surface of the meristem, within the outer cell layer, and is neighboured by its daughter cells which together form the stem cell niche (Fig. 1). To identify genes that are expressed in these discrete domains within the mature *M. polymorpha* meristem, we generated translational and transcriptional reporter lines for three candidate genes based on published literature.

To illustrate the asymmetry of stem cell niche position within the dorsoventral axis of the mature meristem, we imaged the mature meristems of Class III Homeodomain-Leucine-Zipper (C3HDZ) protein fusion lines reported in Wallner and Dolan, 2024 (Fig. 2A-A’’, Fig. S1A-A’’). C3HDZ proteins are required for meristem formation, meristem maintenance, and adaxial-abaxial specification of leaf primordia in *Arabidopsis thaliana* (McConnell et al., 2001; Talbert et al., 1995), and MpC3HDZ localises to the dorsal meristem surface of *M. polymorpha* during meristem formation in the sporeling (Wallner & Dolan, 2024). In the sagittal plane of plants expressing *pro*Mp*C3HDZ*:*g*Mp*C3HDZ*-*VENUS* in the mature meristem, the MpC3HDZ-VENUS signal was restricted to the dorsal surface of the meristem, in the cell layers forming the air chambers (Fig. 2A’). Expression was detected in daughter cells formed from a dorsal division of the apical cell, and not in daughter cells derived from a ventral division of the apical cell. In the frontal plane, VENUS signal was weaker than the dorsal signal in the sagittal plane (Fig. 2A). Similarly, signal was low in the transverse plane and was only detectable above the apical cell (Fig. 2A’’). We conclude that Mp*C3HDZ* reporter expression was enriched in the dorsal surface of the mature *M. polymorpha* meristem.

**Figure 2:**
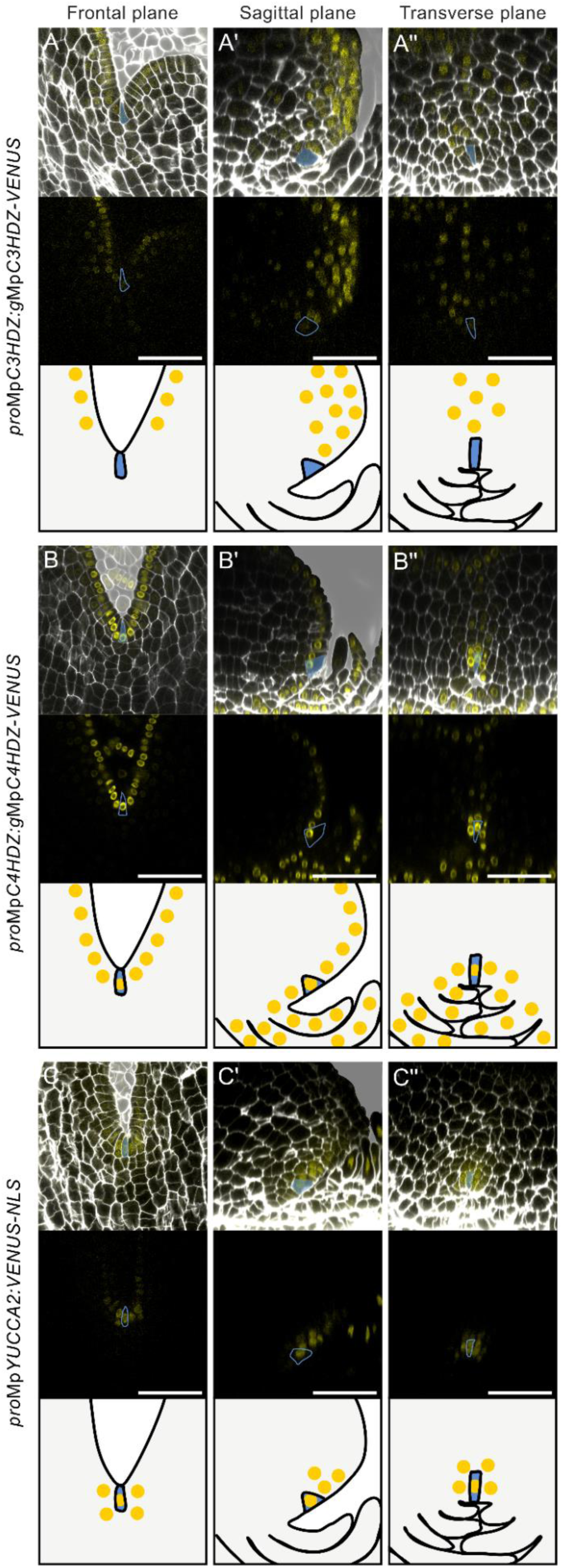
Reporter expression is distinct in each anatomical plane of the mature meristem and is asymmetric in the dorsoventral and apical-basal axes. (**A**-**C’’**) Meristems of reporter lines in the Tak-1 x Tak-2 background grown for four weeks in white light. Lines include (**A**-**A’’**) *pro*Mp*C3HDZ*:*g*Mp*C3HDZ*-*VENUS*, (**B**-**B’’**) *pro*Mp*C4HDZ*:*g*Mp*C4HDZ*-*VENUS*, and (**C**-**C’’**) *pro*Mp*YUC2*:*VENUS*-*NLS*. SR 2200 cell wall stain is shown in white, and VENUS signal is shown in yellow. In the top panels, the space surrounding the meristem is marked in pale grey for clarity. In the middle panels, VENUS signal alone is shown. In the bottom panels, a schematic representation of the expression pattern is shown. The predicted apical cell is marked in blue throughout. n=10, n=10, n=7 for A-A’’, B-B’’ and C-C’’ respectively. All samples showed consistent expression patterns. (**A, B, C**) Frontal plane of the meristem. (**A’**, **B’**, **C’**) Optical reconstruction of the sagittal plane of the meristems in A, B, C. (**A’’**, **B’’**, **C’’**) Optical reconstruction of the transverse plane of the meristems in A, B, C. A second independent line for each reporter is shown in Supplementary Figure 1. Scale bars: A-C’’= 50 µm.

To visualise the outer cell layer, we generated Class IV HDZIP (C4HDZ) protein fusion lines (Fig. 2B-B’’, Fig. S1B-B’’), as C4HDZs are required for epidermal development in *A. thaliana* and some family members such as At*ATLM1* and At*PDF2* are expressed in the L1 layer of the meristem (Abe et al., 2003; Nagata & Abe, 2023). In plants transformed with *pro*Mp*C4HDZ*:*g*Mp*C4HDZ*-*VENUS*, MpC4HDZ-VENUS signal was restricted to the outer cell layer in all three planes of the mature *M. polymorpha* meristem (Fig. 2B-B’’). Within the outer cell layer, we observed stronger signal in the stem cell niche than in neighbouring cells (Fig. 2B-B’’). MpC4HDZ-VENUS signal was also detected in scales which were one cell layer thick and developed from epidermal cells. MpC4HDZ-VENUS signal therefore marked the outer cell layer, including the stem cell niche.

To visualise the stem cell niche, we generated *YUCCA* (*YUC*) transcriptional reporter lines (Fig. 2C-C’’, Fig. S1C-C’’). *YUC* is a conserved auxin biosynthesis gene family in land plants and YUC proteins localise to sites of high auxin production, including the root apical meristem and shoot apical meristem of flowering plants and the meristem of *M. polymorpha* (Blakeslee et al., 2019; Eklund et al., 2015; Hirakawa et al., 2020). In the frontal plane, *pro*Mp*YUC2*:*VENUS*-*NLS* reporter signal was high in the stem cell niche and neighbouring cells (Fig. 2C). The sagittal plane revealed that the strongest signal was detected in the dorsal region above the stem cell niche at the thallus apex (Fig. 2C’). We conclude that *pro*Mp*YUC2*:*VENUS*-*NLS* is highly expressed in a restricted region in and above the stem cell niche.

In summary, the VENUS signals of marker lines for MpC3HDZ, MpC4HDZ and Mp*YUC2* were spatially restricted to the dorsal surface, the outer cell layer (including the apical cell), and the stem cell niche of the mature meristem, respectively. Consistent with meristem anatomy (Fig. 1), reporter signal was asymmetric in the dorsoventral and apical-basal axes, but symmetric in the left-right axis. This expression asymmetry was only evident when examined in the frontal, sagittal and transverse planes.

Mature *M. polymorpha* meristems are derived from meristems that develop on sporelings, or on small vegetive propagules known as gemmae. Gemmae form inside gemma cups on the dorsal surface of the thallus, where they remain dormant. After dispersal, the gemmae meristems are activated and gemmaling growth begins (Kato et al., 2020). It is unclear how the architecture of the meristem changes during gemmaling development and how this relates to the mature meristem. Therefore, we characterised the structure of the meristem during gemmaling development (Fig. 3).

**Figure 3:**
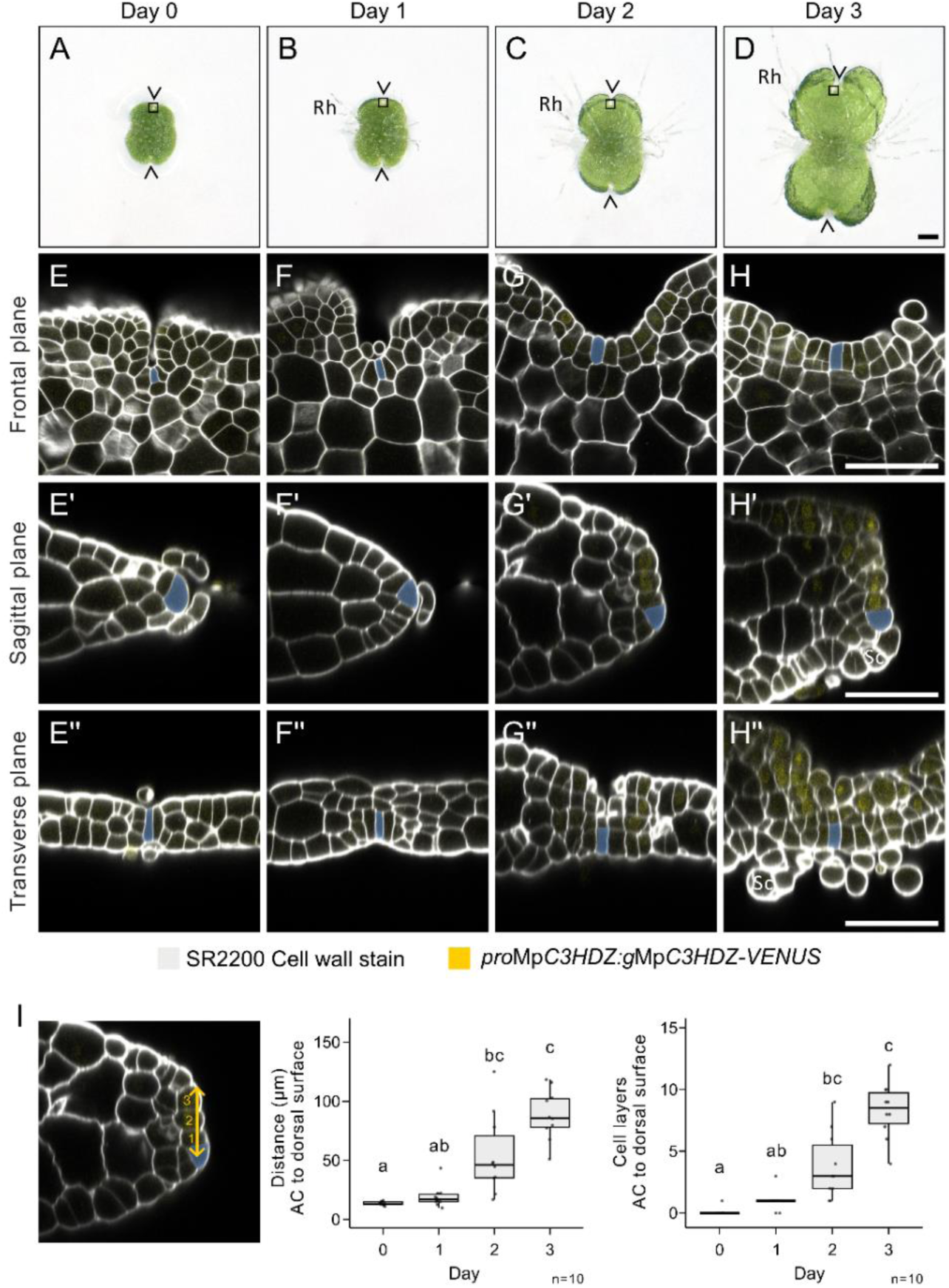
Dorsoventral anatomy and asymmetry is established from a symmetrical gemma meristem. (**A-D**) Keyence timelapse images of a Tak-1 xTak-2 *pro*Mp*C3HDZ*:*g*Mp*C3HDZ*-*VENUS M. polymorpha* gemmae for 3 days following transfer from gemma cup to media. (**A**) On day 0, immediately after transfer from the gemma cup, there are 2 notches on the gemma (arrowheads). (**B**) After 1 day, rhizoids (Rh) are formed from the dorsal and ventral surface. (**C**) On day 2, gemmaling size increases. (**D**) On day 3, notch depth increases, and new rhizoids are only initiated from the ventral surface. Squares show the typical position of the meristems imaged and measured in E-I. (**E**-**H**) Confocal microscopy optical sections in the frontal plane on day 0-3. The predicted apical cell (blue) is located at the base of the notch in the outer cell layer at all time points. (**E’**-**H’**) A reconstruction of the sagittal plane of E-H. The apical cell (blue) progressively moves from the apex (E’ and F’) to the ventral side of the thallus (G’ and H’). The thallus thickness increases, and ventral scales (Sc) form at day 3 (H’). (**E’’**-**H’’**) A reconstruction of the transverse plane of E-H. The apical cell (blue) is located in the centre of the dorsoventral axis at day 0 (E’’), but then localises to the ventral side of the thallus as the total thallus thickness increases (F’’-H’’). Scales (Sc) are evident in H’’. n= 10, n= 6, n=12 and n=7 for E-E’’, F-F’’, G-G’’ and H-H’’ respectively. 0/10, 2/6, 10/12 and 7/10 samples showed reporter signal at each time point. (**I**) Quantification of the distance (yellow line) and the number of cell layers (yellow numbers) from the predicted apical cell (AC, blue) to the dorsal meristem surface. Fig. 2G’ is shown in panel I to explain how the distance and cell layer number were calculated. Kruskal-Wallis tests with Dunn’s multiple comparisons were performed [χ^2^(3)=29.65, p<0.0001; χ^2^(3)=32.35, p<0.0001]. Lower case letters denote statistical difference with a p-value of <0.05. n= 10 for each time point. The transgenic line used was the same as in Fig. S1A-A’’. Scale bars: A-D= 200 µm; E-H’’= 50 µm.

Gemmae meristems were fixed immediately from the gemma cup (Day 0), and one, two and three days after transfer to media (Fig. 3A-D). In the frontal plane, the stem cell niche was located at the base of the notch at all time points (Fig. 3E-H). The apical cell was rectangular and located in the outer cell layer, which formed a U-shaped invagination. The notch width expanded from day 2 as the meristem started to bifurcate (Fig. 3G-H, Fig. S2A-C). The tissue morphology on the left and right of the meristem was identical on each day.

In the sagittal plane, the ungerminated gemma (Day 0) was four cell layers thick with a subtending slime papilla above and below the apical cell (Fig. 3E’). The apical cell was positioned at the tip (apex) and the morphology of the tissue above the apical cell was identical to the tissue below the apical cell. 6/10 gemmae had one apical cell, whilst 4/10 gemmae had two apical cells within the same meristem, both equally sized and positioned at the apex (Fig. S2D-E). After 2 days, the stem cell niche was asymmetrically localised to one side of the apex (Fig. 3G’-H’).

To confirm that the stem cell niche was displaced to the ventral side of the apex, we examined the expression pattern of the *pro*Mp*C3HDZ*:*g*Mp*C3HDZ*-*VENUS* reporter, which marks the dorsal surface of the mature meristem (Fig. 2A-A’’). VENUS signal was undetectable at day 0 and day 1 when the apical cell was positioned at the apex, but was present in cells above the apical cell at day 2 (Fig. 3E’-H’). The stem cell niche was therefore positioned on the ventral side of the gemmaling from day 2 (Fig. 3G’). Both the absolute distance and the number of cell layers between the apical cell and the dorsal meristem surface increased during day 1-3 (Fig. 3I).

In the transverse plane, the lobes to the left and right of the gemma meristem were two cell layers thick, and there were slime papillae above and below the apical cell (Fig. 3E’’). The morphology of the tissue to the left and right of the apical cell was identical, and the morphology of the tissue above and below the apical cell was identical. During the next two days, more tissue layers were produced, and the stem cell niche was localised to the ventral side of the gemmaling (Fig. 3G’’-H’’). The ventral position of the stem cell niche was supported by dorsal expression of *pro*Mp*C3HDZ*:*g*Mp*C3HDZ*-*VENUS*. Furthermore, ventral structures such as scales were evident three days after removal from the gemma cup (Fig. 3H’).

These data indicate that the day 0 gemma meristem has left-right symmetry that persists through gemmaling development and into the mature meristem. However, unlike the mature meristem, the day 0 gemma meristem has dorsoventral symmetry. Anatomical dorsoventral asymmetry is detectable from day 2 and coincides with increased tissue layer formation and Mp*C3HDZ* reporter expression. We conclude that the dorsoventral anatomy and asymmetry that define the mature meristem are established during gemmaling growth.

During the first 3 days of gemmaling development, the stem cell niche becomes displaced to the ventral surface and dorsoventral asymmetry is established (Fig. 3). However, the anatomy of the gemmaling meristem at day 3 differs from the mature meristem at week 4. To determine if the overall anatomy of the meristem and the position of the stem cell niche stabilises in the mature plant, fixed meristem samples were cleared and imaged one week, two weeks, three weeks, and four weeks after gemmae transfer to media (Fig. 4A-D).

**Figure 4:**
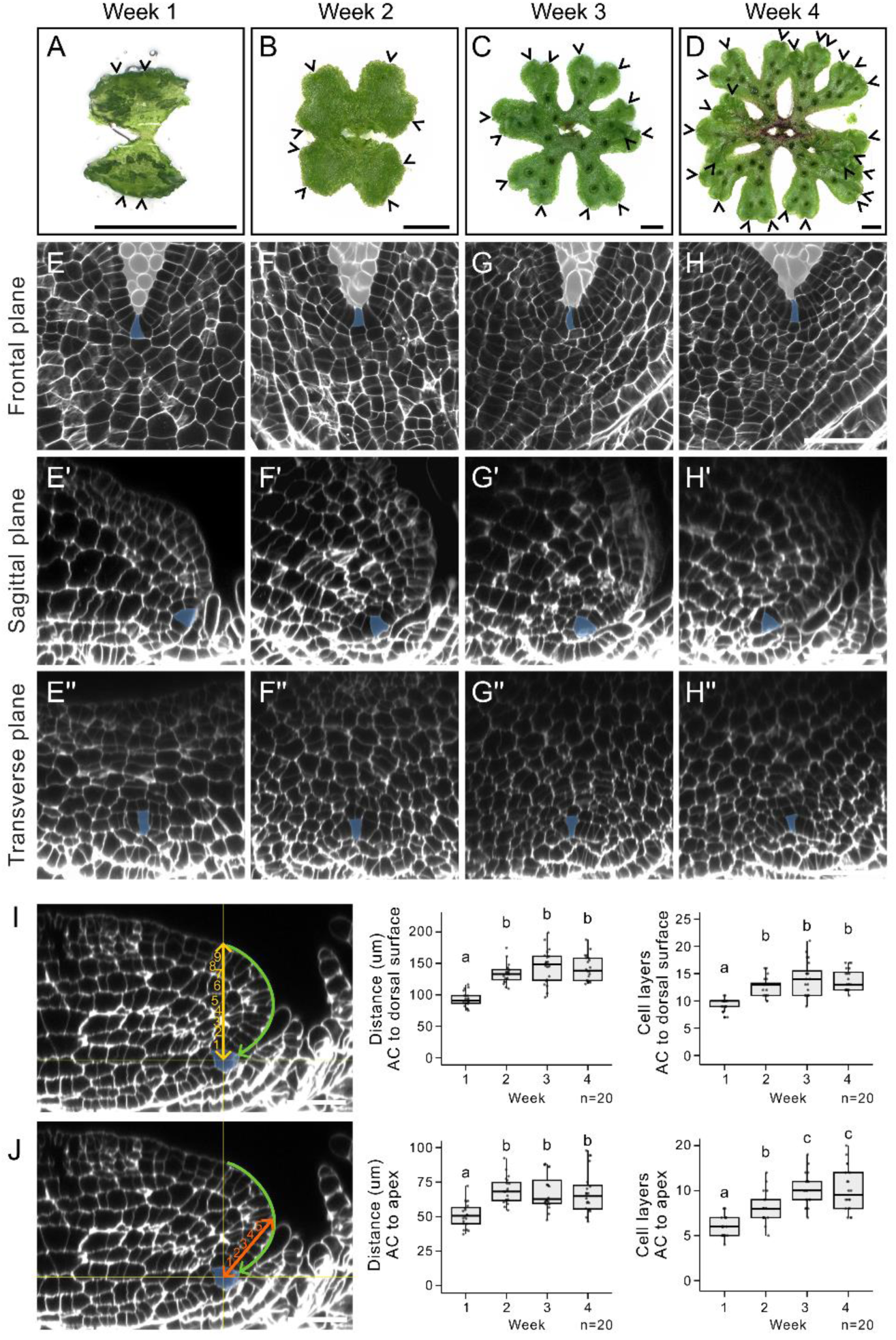
The stem cell niche is maintained in a sub-apical and ventral position after 1 week of growth. (**A**-**D**) Keyence images of Tak-1 *Marchantia polymorpha* plants grown for (**A**) 1, (**B**) 2, (**C**) 3 and (**D**) 4 weeks in white light following removal from the gemma cup. (**E**-**H**) Frontal plane of meristems. The predicted apical cells (blue) are located at the base of the notch and meristem architecture is unchanged during this period. (**E’**-**H’**) Optical reconstructions of the sagittal plane of E-H. The apical cells (blue) are located on the ventral side of the thallus body, behind the protruding thallus apex. (**E’’**-**H’’**) Optical reconstructions of the transverse plane of E-H. The apical cells (blue) are located on the ventral side of the thallus and are surrounded by scales throughout. The space surrounding the meristem is marked in pale grey for clarity in E-H. The experiment was repeated two times with consistent results. (**I**) The vertical distance (yellow line) and the number of cell layers (yellow numbers) between the predicted apical cell (AC, blue) and the dorsal surface were measured. Welch’s ANOVA tests with Games-Howell multiple comparisons were performed [F(3,40.61)=51.96, p<0.0001; F(3, 39.58)=40.7, p<0.0001]. n= 20 for each time point. (**J**) The distance (orange line) and number of cell layers (orange numbers) between the predicted apical cell and the thallus apex were measured. Kruskal-Wallis tests with Dunn’s multiple comparisons were performed [χ^2^(3)=25.74, p<0.0001; χ^2^(3)=41.03, p<0.0001]. n= 20 for each time point. Lower case letters denote statistical difference with a p-value of <0.05. Scale bars: A-D= 5 mm; E-J= 50 µm.

In the frontal plane, the apical cell was located at the base of the notch throughout development (Fig. 4E-H). The apical cell was trapezoid and in the outer cell layer at each time point. We did not detect any differences in meristem anatomy in this plane between week 1 and 4. In the sagittal plane, the stem cell niche was ventrally and sub-apically localised and was surrounded by scales throughout the four-week time course (Fig. 4E’-H’). In the transverse plane, the stem cell niche was ventrally localised and surrounded by scales from week 1 to 4 (Fig. 4E’’-H’’). Therefore, the symmetry and arrangement of the meristem was consistent throughout the four-week time course.

To determine if the relative ventral position of the stem cell niche changed during this time course, the vertical distance between the apical cell and the dorsal surface was measured at each time point (Fig. 4I). After an initial increase from week 1 to 2, there was no change in the total distance or number of cell layers between the apical cell and the dorsal surface from week 2 to 4 (Fig. 4I). To determine if the sub-apical position of the stem cell niche changed with age, the distance between the apical cell and the thallus apex was measured (Fig. 4J). The distance between the apical cell and the thallus apex increased from week 1 to 2 and then remained constant from week 2, whilst the number of cell layers remained constant from week 3 (Fig. 4J).

We conclude that the overall asymmetry and anatomy of the meristem was similar throughout weeks 1 to 4. The stem cell niche was positioned ventrally and sub-apically throughout weeks 1 to 4, however its relative position within the meristem became stable 2-3 weeks after removal from the gemma cup. We conclude that the stem cell niche first becomes displaced to the ventral surface from day 2, before being displaced sub-apically between day 3 to 7.

## DISCUSSION

Meristems maintain a population of stem cells whilst producing cells that differentiate into tissues and organs. This balance of stemness and differentiation is achieved by spatially patterned gene expression networks that operate in the three-dimensional anatomy of the meristem (Eshed Williams, 2021; Kohchi et al., 2021). We report the three-dimensional structure of the mature *Marchantia polymorpha* meristem and show that the spatial localisation of MpC3HDZ, MpC4HDZ and Mp*YUC2* reporter signal is restricted to the dorsal surface, outer cell layer, and stem cell niche of the meristem respectively. The dorsoventral asymmetry of the mature meristem anatomy is consistent with the asymmetry of the reporter signal. We demonstrate how the three-dimensional structure of the meristem changes during vegetative development. The cellular anatomy of the gemma meristem is relatively simple with dorsal-ventral symmetry. To generate the dorsal-ventral asymmetry characteristic of the mature meristem, the stem cell niche is displaced towards the ventral surface from day 2, and then becomes positioned sub-apically between day 3-7. The relative position of the stem cell niche within the meristem is stable after 2 weeks (Fig. 5).

**Figure 5:**
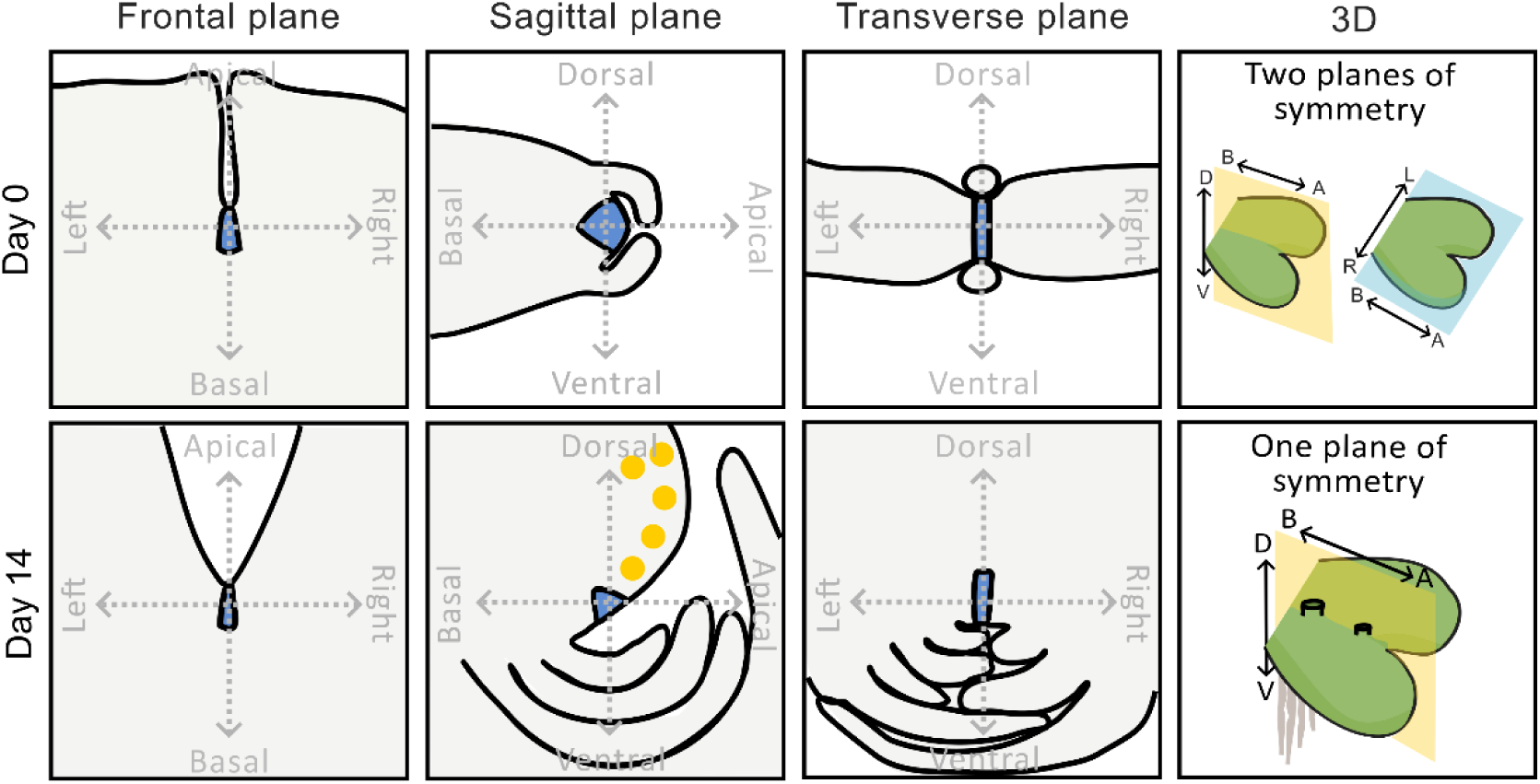
Summary diagram of meristem anatomy in three anatomical planes at day 0 and day 14. At day 0, the gemma meristem is notched-shaped in the frontal plane, with left-right symmetry. In the sagittal plane, the meristem has dorsoventral symmetry. Both the left-right symmetry and the dorsoventral symmetry of the day 0 gemma are evident in the transverse plane. We conclude that there are two planes of symmetry at this time point, which correspond to the sagittal and frontal planes. During the following 14 days, the gemmaling meristem undergoes anatomical rearrangement. The left-right symmetry of the meristem is maintained, however the dorsoventral symmetry is lost. Therefore, unlike the gemma meristem, the mature meristem has only one plane of symmetry in the sagittal plane. Both the anatomy and gene expression profiles of the mature meristem are asymmetrical: Mp*C3HDZ* reporter line signal (yellow) is restricted to the dorsal meristem surface only. We conclude that the gemmae meristem represents an intermediate stage of meristem development, that develops during the following two weeks into the meristem of the mature *M. polymorpha* plant.

The mature *M. polymorpha* meristem has left-right symmetry, dorsoventral asymmetry, and apical-basal asymmetry. The meristem is therefore bilaterally symmetrical, unlike angiosperm shoot and root apical meristems which are often radially symmetrical (Doerner, 2003; Moubayidin, 2015). The dorsoventral asymmetry of the *M. polymorpha* mature meristem is reflected in the asymmetrical expression of regulatory genes. For example, Mp*C3HDZ* reporter expression is low in the frontal plane but is strongly expressed in the dorsal domain of the meristem in the sagittal plane. This is consistent with published data showing that Mp*C3HDZ* regulates dorsoventral identity (Wallner & Dolan, 2024). Furthermore, enrichment of Mp*YUC2* reporter expression in the cells on the dorsal side of the stem cell niche is only evident in the sagittal plane. Previous reports of *pro*Mp*YUC2*:*GUS* expression showed that GUS signal was restricted to the notch, but did not show cellular level resolution or the restriction of expression to the dorsal surface (Eklund et al., 2015; Hirakawa et al., 2020; Takahashi et al., 2021). Our data indicate that the expression patterns of genes within the meristem are asymmetric, highlighting the necessity to examine all three planes to gain a complete picture of gene expression in the three-dimensional meristem. Many GUS reporter lines for developmental genes in Marchantia have been shown to localise to the notch (Aki et al., 2019; Flores-Sandoval et al., 2015, 2018; Fu et al., 2024; Hirakawa et al., 2019, 2020; Liu et al., 2024), and our results provide a framework to identify their three dimensional expression in cellular detail in the meristem.

Our temporal analysis of meristem anatomy from gemmae reveals that the meristem undergoes substantial change during the first three days of gemmaling development. This includes the establishment of dorsoventral asymmetry, which can develop in either orientation depending on which side the gemma lands after dispersal (Bowman, 2016; Mirbel, 1835). Light has been shown to polarise the gemma body; after the gemma lands, the surface towards the light source develops dorsal identity and the surface furthest away develops ventral identity (Bowman, 2016; Mirbel, 1835; Otto & Halbsguth, 1976). We found that Mp*C3HDZ* reporter signal was only detectable in the gemmaling two days after removal from the gemma cup. The timing of MpC3HDZ-VENUS reporter signal and asymmetry establishment correlates with early publications that showed that dorsoventral polarity is irreversible after 1-3 days (Bowman, 2016; Mirbel, 1835; Pfeffer, 1871), and may correspond to light-induced polarity establishment. Changes in localisation of MpC3HDZ-VENUS during gemmaling development are also consistent with changing gene expression patterns during day 0 to day 3 for Mp*ERF20*, Mp*BZIP15* and Mp*C2H2-22* transcriptional reporter lines (Romani et al., 2024), and may be associated with the anatomical maturation of the meristem and the suppression of meristem dormancy. Our data are consistent with the hypothesis that the gemma meristem is an arrested meristem at an early stage of development. Upon removal from the gemma cup, gemma meristem development is re-initiated, leading to the formation of a mature meristem.

Our results provide the first spatio-temporal characterisation of meristem anatomy during the development of the haploid phase of *M. polymorpha*. We reveal that the meristem is itself dorsoventralised and this dorsoventrality is specified by the environment after the establishment of apical-basal and left-right body axes during gemmae development. This contrasts with other meristems such as *A. thaliana,* where the apical meristems are radially symmetrical and dorsoventrality of mature organs develops in primordia and not in the meristem.

## MATERIALS AND METHODS

### Plant material and growth conditions

*Marchantia polymorpha* wild type accessions Takaragaike-1 (Tak-1, male) and Takaragaike-2 (Tak-2 female), and transgenic plants were grown in continuous white light at 45 µmol m^-2^ s^-1^ at 23°C (see supplementary table for spectrum). Plants for figure 1 and 2 were grown for two weeks on plates containing ½ B5 Gamborg’s Media [½ strength B5 Gamborg + 0.5 g/L MES hydrate + 1% sucrose + 1% plant agar, pH 5.5] and then transferred to SacO2 Microboxes containing autoclaved 3:1 compost: vermiculite mix for a further two weeks. Plants in figure 3 were grown for 3 days on media plates, and plants in figure 4 were grown for 4 weeks on media plates without transfer to soil. To generate spores for transformation, two-week-old plants grown on media plates in continuous white light were transferred to soil and moved to 16 hr far-red enriched light (50 µmol m^-2^ s^-1^, see supplementary table for spectrum), 8 hr dark, at 20°C. Plants were crossed and the mature sporangia harvested. Sporangia were dried for 4 weeks before freezing at -70°C.

### Generation of plasmids for transformation

To generate transcriptional and translational reporter constructs, we followed the Greengate protocol (Lampropoulos et al., 2013). The candidate gene promoter drives a nuclear localised fluorescent gene in transcriptional reporters (VENUS-NLS), whilst the candidate gene promoter drives expression of the candidate gene genomic DNA sequence fused to VENUS in the translational fusion reporters. *pro*Mp*C3HDZ*:*g*M*pC3HDZ*-*VENUS* lines were generated in Wallner et al., 2024. Promoter and genomic gene sequences were amplified from genomic Tak-1 DNA by CloneAmp HiFi PCR Premix with Green Gate-specific primers and cloned via BsaI restriction sites into entry modules with a pUC19 based vector backbone. The Mp*YUC*2 promoter was defined as region 4001bp upstream of ATG start codon. The *MpC4HDZ* promoter region was defined by a region 4309bp upstream of the ATG. The genomic sequence of Mp*C4HDZ* encompassed 3627bp (including exons and introns without the stop codon). To generate promoter-reporter *proMpYUC2:VENUS-linker-NLS* and translational fusion *pro*Mp*C4HDZ*:*g*M*pC4HDZ*-*VENUS*, Greengate entry modules listed in the supplementary table were assembled into the pGreen-IIS based destination vector. The linker-VENUS module for the translational fusion constructs was published previously (Wallner et al., 2023) and the chlorsulfuron plant resistance module was adapted from the OpenPlant toolkit (Sauret-Güeto et al., 2020). Cloning was performed by the Protein Technologies Facility, VBC.

### Generation of transgenic lines

Spores generated from a Tak-1 x Tak-2 cross were transformed with reporter constructs according to the protocol described in Wallner et al., 2024, adapted from Ishizaki et al., 2008. Sporelings were plated on ½ B5 Gamborg’s Media containing 0.5 µM Chlorsulfuron and 100 mg/L Cefotaxime. Positive transformants were grown on non-selective plates and screened for reporter expression. At least 10 independent lines were screened, and four lines (2 males and 2 females) were selected that showed consistent and strong reporter signal. Reporter expression in one independent line is shown in Figure 2, and a second independent line is shown in Supplementary Figure 1. The lines were verified using “Plant sequencing primers” listed in the supplementary table. Plants were stored as gemmae stocks in media stabs at 4°C, and stored on media plates as mature plants in continuous white light at 45 µmol m^-2^ s^-1^ at 17°C (see supplementary table for spectrum).

### Tissue fixation and clearing

The notches of plants were harvested and transferred immediately to 10% Formalin Solution (10% Neutral Buffered, 4% w/v formaldehyde) + 0.1% Brij® L23 solution. Samples were vacuumed twice for 25 minutes each, and then washed three times in phosphate buffered saline (PBS). PBS solution was replaced with Clear-See α solution [10% (w/v) Xylitol + 15% (w/v) Sodium deoxycholate + 25% (w/v) Urea + 50 mM Sodium Sulfite Anhydrous]. Samples were vacuumed for 25 minutes, and then incubated in the dark on a shaker overnight in Clear-See α solution. The next day the solution was replaced, and samples were incubated in the dark on a shaker until imaging (minimum 5 days).

### Imaging and Microscopy

Live plants were imaged using the Keyence VHX-7000 digital microscope with the VH-ZST and VH-Z00R/W/T objectives.

For fixed samples, 0.2% SR 2200 cell wall stain was added to cleared samples the day before imaging. Samples were mounted on glass slides in Clear-See α solution in 0.25mm thick Gene Frames, and imaged using an inverted point scanning Zeiss LSM880 confocal microscope with an 40x/1.2 LD LCI plan-apochromat, Water, Glycerol DIC AutoCorr Objective and MBS 458/514 and MBS -405 filters. Silicon Oil was used for immersion and Z-stacks of 300-600 slices were acquired depending on the thickness of the sample. The recommended z-stack interval was selected. To detect SR 2200 signal, a 504 nm diode laser (1-2%) was used for excitation and signal was detected between 419-499 nm with a 32 µm pinhole (∼1 AU). To detect VENUS signal, a 514 nm argon laser (20%) was used for excitation and VENUS signal was detected between 526-579nm with a 39 µm pinhole (∼1 AU). The two channels were scanned sequentially in all imaging. For images shown in figure 2, the track was switched after each frame, whilst for figure 3 the track was switched after each z-stack.

For Fig. 1H, L, week 4 samples stained with SR 2200 were imaged using a Zeiss Z1 Lightsheet microscope. Samples were mounted in 3% Low Melting Agarose in Clear See solution and incubated at 4°C for 1-3 hours. An agarose block containing the sample was cut and glued to a plastic support, which was attached to a capillary tube held within the sample mount. The large sample imaging chamber (n= 1/33-1.58) was filled with Clear See solution. LSFM 5x/0.1 foc excitation objectives were used with a 5x/0.16 (EC-Plan Neofluar 5x/0.16 foc) detection objective. A 405nm 20mW laser (5%) was used to detect SR 2200 signal.

### Image analysis

The Z-stacks acquired were used to reconstruct the sagittal and transverse planes using Orthogonal Views in Fiji. Stacks were rotated to produce sagittal planes through the geometric centre of the notch. For figure 3 and 4, the sagittal plane was measured at all stages of the bifurcation cycle. The distance between the apical cell and the thallus apex, and the distance between the apical cell and the dorsal surface was measured. Cell layers were counted along these measurement axes; a cell layer was counted each time a new cell wall was crossed. Meristems that had recently produced gemma cups were excluded from the analysis. All meristems shown in figures had recently bifurcated and the central lobe had expanded. This stage was chosen because the next round of bifurcation (and thus stem cell duplication) was least likely to be occurring. Experiments were performed once unless stated otherwise in the figure legend.

### Statistical analysis and data presentation

All statistical analysis was performed using R-studio. Welch’s ANOVA tests with Games-Howell multiple comparisons were used when data was normally distributed with unequal variance. When data was not normally distributed, Kruskal-Wallis with Dunn’s multiple comparisons tests were used. All figures were made using Inkscape.

### Reagents, equipment and software

Further manufacturer and identification details for all consumables and equipment can be found in the Supplementary table.

## ACKNOWLEDGEMENTS

We would like to thank the Protein Technologies Facility, the BioOptics Facility, and the Plant Sciences Facility at the Vienna BioCenter for their extensive support and advice. We thank Hugh Mulvey for developing the Clear See protocol in our laboratory and providing valuable feedback on the manuscript, and Zohar Meir for thoughtful discussion. We also thank the Media Kitchen, Lab Support and the administrative staff at the VBC.

## DATA AND RESOURCE AVAILABILITY

All relevant data and resource can be found within the article and its supplementary information.

## AUTHOR CONTRIBUTIONS

V.S. and L.D. designed the project. V.S. carried out the project. E.-S.W. designed the constructs. K.J. and V.S. potted out plants, harvested samples and prepared microscopy slides. E.-S.W., N.E and V.S. screened reporter lines and N.E. maintained plant stocks. M.M. transformed spores to generate reporter lines. V.S. and L.D. wrote the manuscript.

## SUPPLEMENTARY FIGURES

**Supplementary Figure 1:**
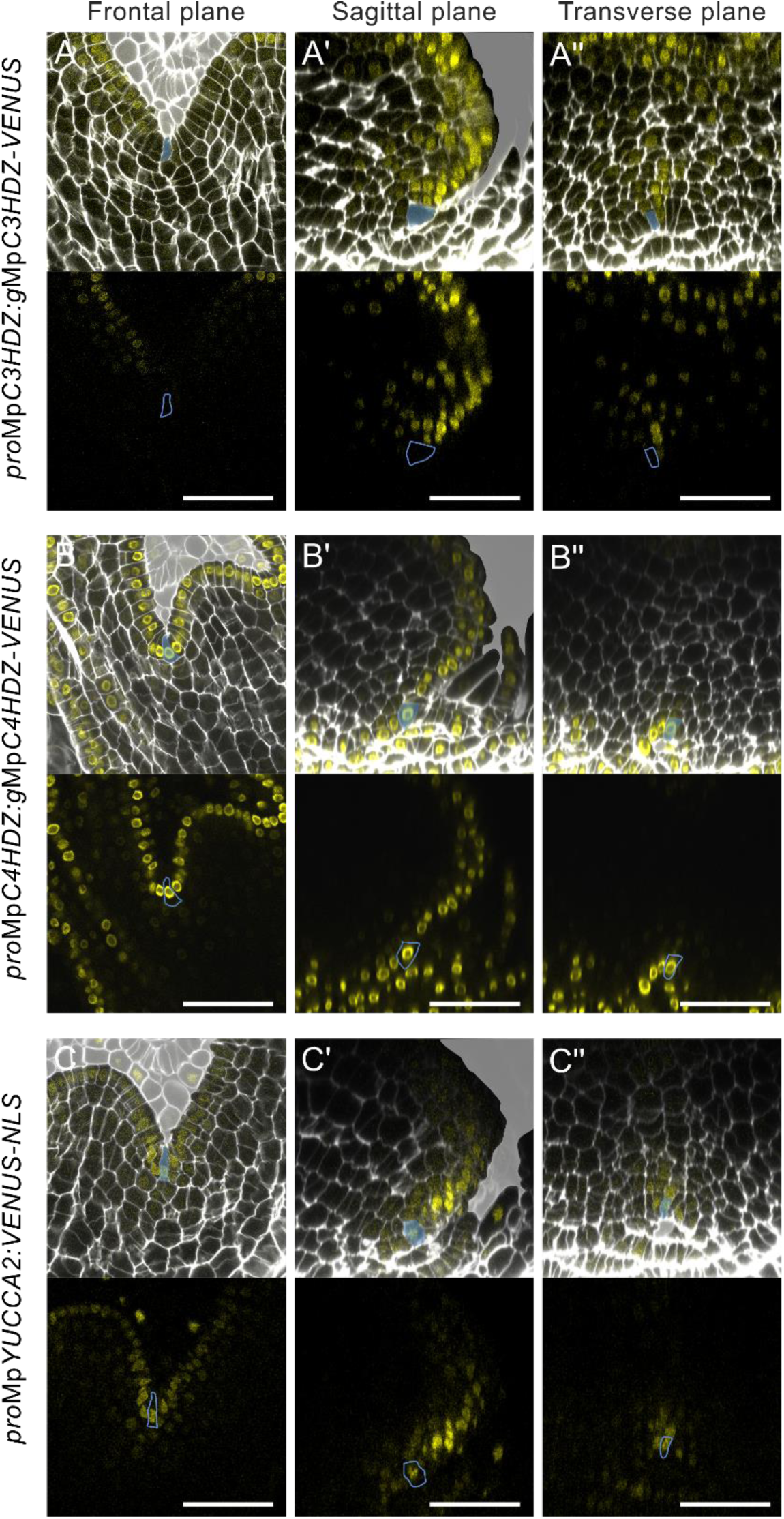
MpC3HDZ, MpC4HDZ and Mp*YUC2* reporter line expression. (**A**-**C’’**) Meristems of a second independent reporter line in the Tak-1 x Tak-2 background grown for four weeks in white light. Lines show the same expression patterns as Figure 2 for (**A**-**A’’**) *pro*Mp*C3HDZ*:*g*Mp*C3HDZ*-*VENUS*, (**B**-**B’’**) *pro*Mp*C4HDZ*:*g*Mp*C4HDZ*-*VENUS*, and (**C**-**C’’**) *pro*Mp*YUC2*:*VENUS*-*NLS*. SR 2200 cell wall stain is shown in white, and VENUS signal is shown in yellow. In the top panels, the space surrounding the meristem is marked in pale grey for clarity. In the bottom panels, VENUS signal alone is shown. The predicted apical cell is marked in blue throughout. n=3, n=9, n=6 for A-A’’, B-B’’ and C-C’’ respectively. All samples showed consistent expression patterns. (**A, B, C**) Frontal plane of the meristem. (**A’**, **B’**, **C’**) Optical reconstruction of the sagittal plane of the meristems in A, B, C. (**A’’**, **B’’**, **C’’**) Optical reconstruction of the transverse plane of the meristems in A, B, C. Scale bars: A-C’’= 50 µm.

**Supplementary Figure 2:**
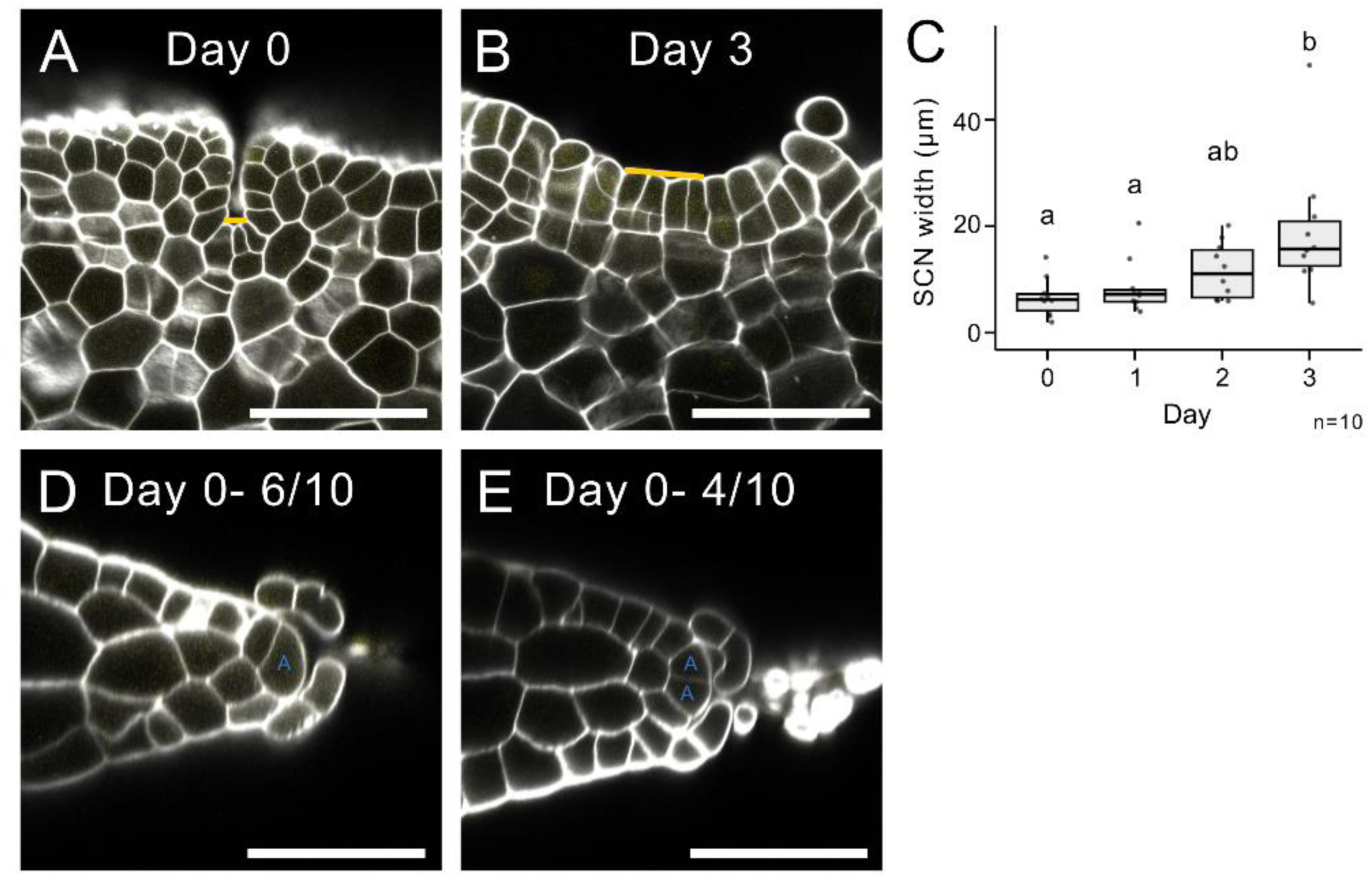
Stem cell niche width increases during day 0-3 of gemmaling development. (**A**-**C**) Measurements of the stem cell niche width, as shown with a yellow line. Cells were included if they were located in the notch base, were smaller than the surrounding cells, and trapezoid/ rectangular in shape. Panels A and B are the images in Fig. 3E and 3H respectively. A Kruskal-Wallis test with Dunn’s multiple comparison was performed [χ^2^(3)=13.15, p=0.0043]. Lower case letters denote statistical difference with a p-value of <0.05. n= 10 for each time point. (**D**-**E**) Day 0 gemma meristems had either one (D) or two (E) detectable apical cells (blue A). 4/10 samples had two apical cells, that were equal in size and both located at the apex. Panel D is the image in Fig. 3E’. Scale bars: A-B, D-E= 50 µm.

### Supplementary Tables

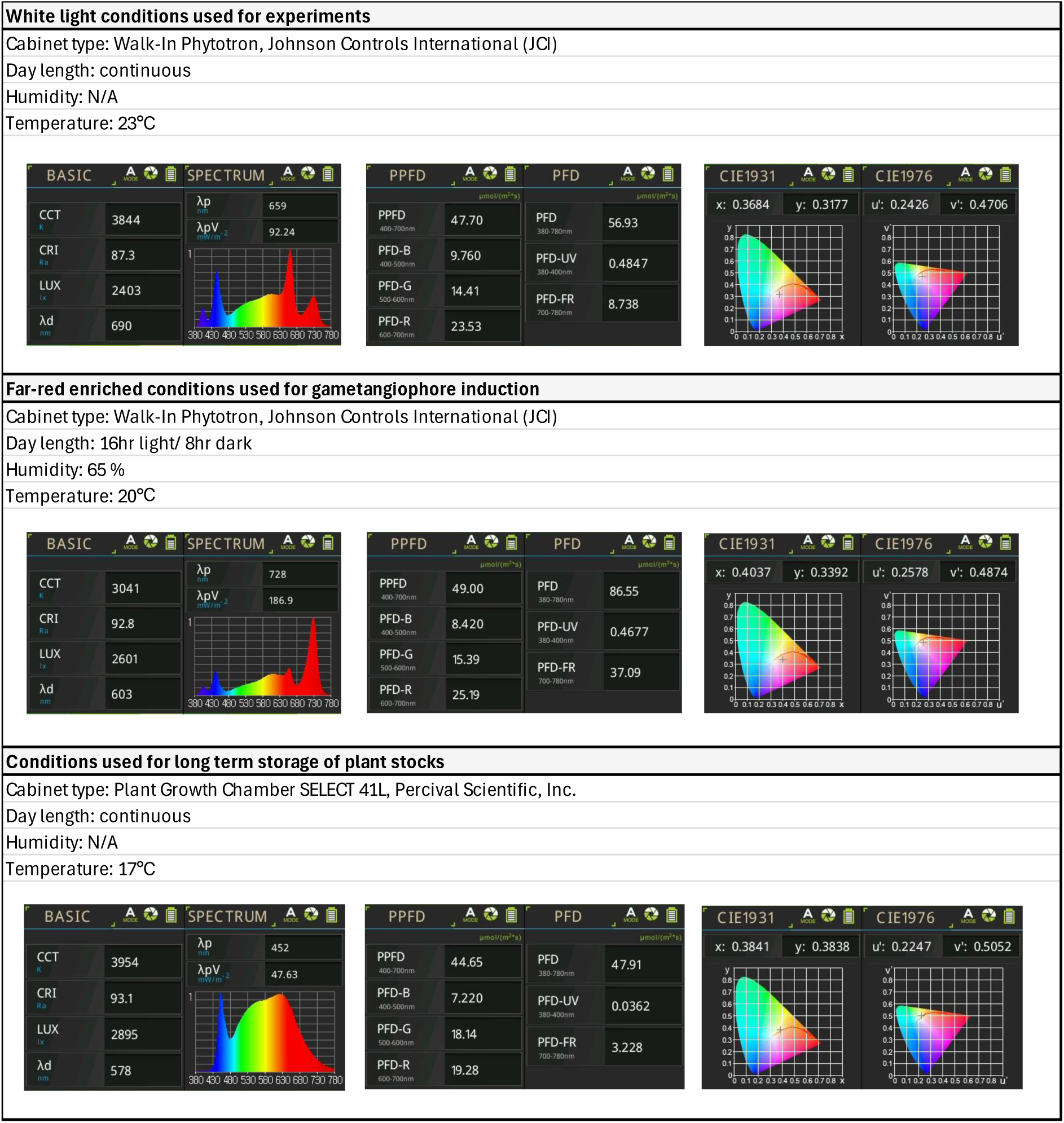

| REAGENT or RESOURCE | SOURCE | IDENTIFIER |
| --- | --- | --- |
| **Consumables:** |  |  |
| Gamborg B5 Medium basal salt mixture | Duchefa | Cat#G0209 |
| MES Monohydrate | Duchefa | Cat#M1503 |
| Sucrose | AppliChem | Cat#A4734 |
| Plant agar | Duchefa | Cat#P1001 |
| Potassium hydroxide (KOH) | Merck | Cat#1.05033 |
| Microbox polypropylene containers, OV80+OVD80 #40 NG/NP | Sac O2, Deinze, Belgium | N/A |
| Neuhaus N3 Compost | Neuhaus | N/A |
| Vermiculite, 2-3mm | Gramoflor | Cat#154618 |
| Chlorsulfuron | Sigma Aldrich | Cat#34322 |
| Cefotaxime sodium | Duchefa | Cat#C0111 |
| Formalin solution, neutral buffered, 10% | Sigma-Aldrich | Cat#HT5012 |
| Brij® L23 solution | Sigma | Cat#B4184 |
| Xylitol | Sigma-Aldrich | Cat#X3375 |
| Urea | AppliChem | Cat#A1049 |
| Sodium deoxycholate | TCI | Cat#C0316 |
| Sodium sulfite anhydrous | Fluka | Cat#71989 |
| Sodium Chloride (NaCl) | Merck | Cat#1.06404 |
| Potassium Chloride (KCl) | Sigma-Aldrich | Cat#60130 |
| Sodium phosphate dibasic (Na<sub>2</sub>HPO<sub>4</sub>) | Sigma-Aldrich | Cat#S9763 |
| Potassium phosphate monobasic (KH<sub>2</sub>PO<sub>4</sub>) | Sigma-Aldrich | Cat#60220 |
| SR 2200 stain | Renaissance Chemicals | N/A |
| Gene frame® 65 µl, 1.5x 1.6 cm | Thermo Scientific | Cat#AB-0577 |
| Gene frame® 125 µl, 1.7x 2.8 cm | Thermo Scientific | Cat#AB-0578 |
| Silicone Immersion Oil | Olympus | Cat#300CS-30CC |
| Agarose Low Melt | Roth | Cat#6351.5 |
| CloneAmp HiFi PCR Premix | Takara | Cat#639298 |
| Sodium dichloroisocyanurate (NaDCC) | Sigma Aldrich | Cat#218928 |
| Acetosyringone | Sigma Aldrich | Cat#D134406 |
| **Equipment:** |  |  |
| Diaphragm Vacuum pump, MZ 2C | Vacuubrand, Wertheim, Germany | N/A |
| Keyence VHX-7000 | Keyence, Osaka, Japan | N/A |
| VH-ZST Objective Lens | Keyence, Osaka, Japan | N/A |
| VH-Z00R/W/T Objective Lens | Keyence, Osaka, Japan | N/A |
| LSM880 Inverted Confocal microscope | Carl Zeiss, Jena, Germany | N/A |
| 40x/1.2 LD LCI plan-apochromat, Water, Glycerol DIC AutoCorr Objective | Carl Zeiss, Jena, Germany | N/A |
| Zeiss Z1 Lightsheet microscope | Carl Zeiss, Jena, Germany | N/A |
| LSFM 5x/0.1 foc excitation objectives | Carl Zeiss, Jena, Germany | N/A |
| EC-Plan Neofluar 5x/0.16 foc detection objective | Carl Zeiss, Jena, Germany | N/A |
| **Software:** |  |  |
| ImageJ/Fiji version 1.54j | www.fiji.sc |  |
| R-studio version 2024.04.2 Build 764 + R version 4.2.2 | www.posit.co |  |
| ZEN 2.3 SP1 FP3 (black) version 14.0.0.0 | www.zeiss.com |  |
| Microsoft 365 MSO version 2401 | www.office.com |  |
| Snapgene version 6.1.2 | www.snapgene.com |  |
| Inkscape 1.3.2 (091e20e, 2023-11-25, custom) | www.inkscape.org |  |

| REAGENT or RESOURCE | SOURCE |
| --- | --- |
| **Plant material:** |  |
| *Marchantia polymorpha* Tak-1 | Ishizaki et al. 2008 |
| *Marchantia polymorpha* Tak-2 | Ishizaki et al. 2008 |
| Tak-1xTak-2 + *pro* MpC3HDZ:g MpC3HDZ -VENUS Line 6f | This paper |
| Tak-1xTak-2 + *pro* MpC3HDZ:g MpC3HDZ -VENUS Line 19m | This paper |
| Tak-1xTak-2 + *pro* MpC4HDZ:g MpC4HDZ -VENUS Line 11f | This paper |
| Tak-1xTak-2 + *pro* MpC4HDZ:g MpC4HDZ -VENUS Line 5m | This paper |
| Tak-1xTak-2 + *pro* MpYUCCA2:VENUS -NLS Line 30f | This paper |
| Tak-1xTak-2 + *pro* MpYUCCA2:VENUS -NLS Line 6m | This paper |
| **Bacterial strains:** |  |
| *Agrobacterium tumefaciens* GV3101: GentR, RifR with pSoup plasmid (TetR) | Made in VBC |
| *Escherichia coli* : strain DH5a | Made in VBC |
| *Escherichia coli* : GreenGate Cloning System | Lampropoulos et al. 2013 |
| **Assembled plasmids:** |  |
| *pro* MpC3HDZ:g MpC3HDZ -VENUS | Wallner et al., 2024 |
| *pro* MpC4HDZ:g MpC4HDZ -VENUS | This paper |
| *pro* MpYUCCA2:VENUS -NLS | This paper |
| **Greengate module plasmids:** |  |
| Greengate backbone- pGGZ003 |  |
| Module A- *pro* MpC3HDZ (4989bp upstream of ATG) | Wallner et al., 2024 |
| Module A- *pro* MpC4HDZ (4309bp upstream of ATG) | This paper |
| Module A- *pro* MpYUCCA2 (4001bp upstream of ATG) | This paper |
| Module B- B-dummy | Lampropoulos et al. 2013 |
| Module C- *g* MpC3HDZ (5756 bp exons + introns) | Wallner et al., 2024 |
| Module C- *g* MpC4HDZ (3627 bp exons + introns) | This paper |
| Module C- VENUS, pGGC000 | Lampropoulos et al. 2013 |
| Module D- Linker VENUS, pGGD000 | Wallner et al., 2023 |
| Module D- Linker NLS | Lampropoulos et al. 2013 |
| Module E- RBCS terminator (from pea) | Lampropoulos et al. 2013 |
| Module F- Chlorsulfuron resistance gene | Sauret-Güeto et al., 2020 |
| **Marchantia.info Gene Identifiers:** |  |
| MpC3HDZ= Mp1g24140 |  |
| MpC4HDZ= Mp7g09710 |  |
| MpYUCCA2= Mp8g08780 |  |

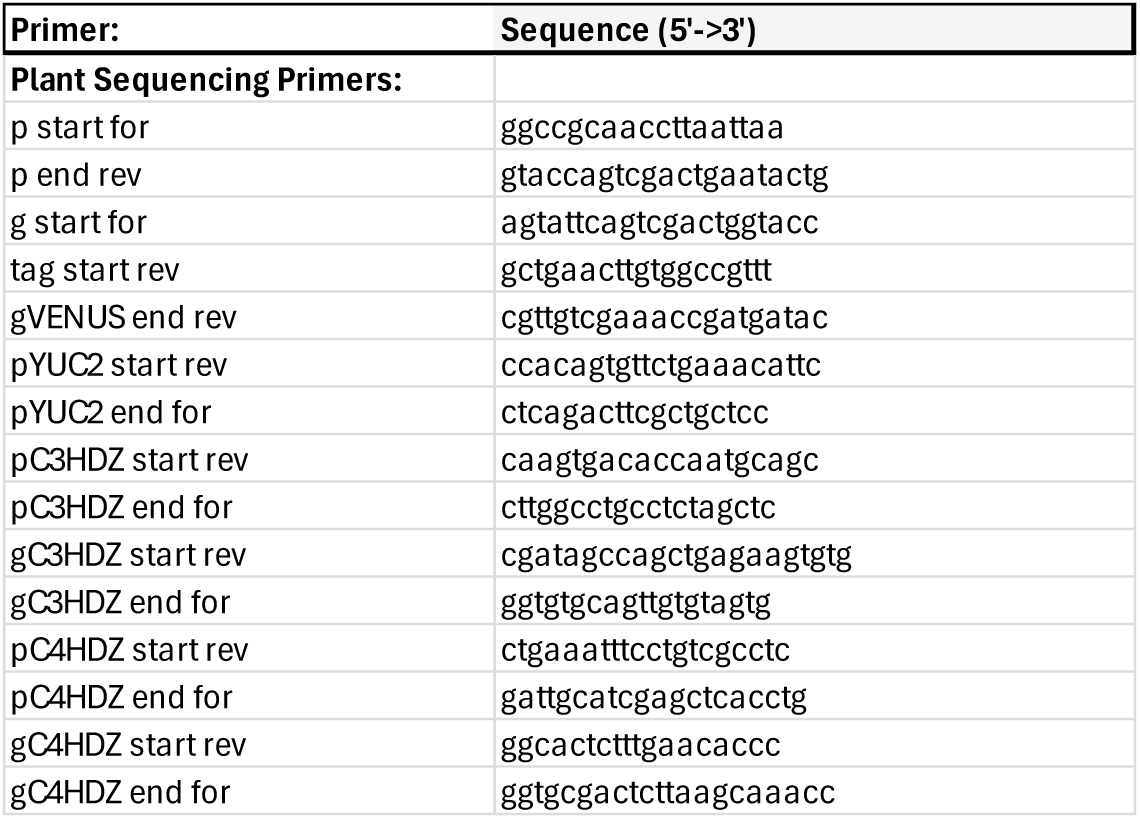

